# Sorcin stimulates Activation Transcription Factor 6α (ATF6) transcriptional activity

**DOI:** 10.1101/2020.04.23.053322

**Authors:** Steven Parks, Tian Gao, Natalia Jimenez Awuapura, Joseph Ayathamattam, Pauline L. Chabosseau, Piero Marchetti, Paul Johnson, Domenico Bosco, Dhananjaya V. Kalvakolanu, Héctor H. Valdivia, Guy A. Rutter, Isabelle Leclerc

**Author notes:** Both authors contributed equally to this work (Steven Parks, Tian Gao). Department of Molecular Mechanisms of Disease, University of Zurich, Switzerland (Tian Gao). Corresponding author: Dr Isabelle Leclerc.

## Abstract

Levels of the transcription factor ATF6α, a key mediator of the unfolded protein response, that provides cellular protection during the progression endoplasmic reticulum (ER) stress, are markedly reduced in the pancreatic islet of patients with type 2 diabetes and in rodent models of the disease, including ob/ob and high fat-fed mice. Sorcin (gene name *SRI*) is a calcium (Ca^2+^) binding protein involved in maintaining ER Ca^2+^ homeostasis.

We have previously shown that overexpressing sorcin under the rat insulin promoter in transgenic mice was protective against high fat diet-induced pancreatic beta cell dysfunction, namely preserving intracellular Ca^2+^ homeostasis and glucose-stimulated insulin secretion during lipotoxic stress. Additionally, sorcin overexpression was apparently activating ATF6 signalling in MIN6 cells despite lowering ER stress.

Here, in order to investigate further the relationship between sorcin and ATF6, we describe changes in sorcin expression during ER and lipotoxic stress and changes in ATF6 signalling after sorcin overexpression or inactivation, both in excitable and non-excitable cells.

Sorcin mRNA levels were significantly increased in response to the ER stress-inducing agents thapsigargin and tunicamycin, but not by palmitate. On the contrary, palmitate caused a significant decrease in sorcin expression as assessed by both qRT-PCR and Western blotting despite inducing ER stress. Moreover, palmitate prevented the increase in sorcin expression induced by thapsigargin. In addition, sorcin overexpression significantly increased ATF6 transcriptional activity, whereas sorcin inactivation decreased ATF6 signalling. Finally, sorcin overexpression increased levels of ATF6 immunoreactivity and FRET imaging experiments following ER stress induction by thapsigargin showed a direct sorcin-ATF6 interaction.

Altogether, our data suggest that sorcin down-regulation during lipotoxicity may prevent full ATF6 activation and a normal UPR during the progression of obesity and insulin resistance, contributing to beta cell failure and type 2 diabetes.

## INTRODUCTION

Obesity and its metabolic consequences are becoming a major threat to human health all over the world, causing life expectancy to fall for the first time in centuries (1). Physical inactivity and nutrient oversupply increase circulating levels of glucose and free fatty acids (FFA), triggering the unfolded protein response (UPR) and endoplasmic reticulum (ER) stress through intensification of the cellular biosynthetic pathways in the pancreatic beta cell (2,3), but also in the liver (4), adipose tissue (5), skeletal muscle (6) and hypothalamus (7). ER stress is also implicated in the development of diabetes complications (8). However, the molecular mechanisms linking glucolipotoxicity and ER stress are not fully understood.

Sorcin (soluble resistance-related calcium binding protein, gene name *SRI,* also known as CP-22, CP22, SCN and V19), is a member of the penta-EF-hand family of calcium binding proteins and relocates from the cytoplasm to membranes in response to elevated calcium levels (9–15). Sorcin was initially identified in multidrug-resistant cancer cells, is ubiquitously expressed and is highly conserved amongst mammals (12,16). In cardiac myocytes and skeletal muscle cells, sorcin inhibits ryanodine receptor (RyR) activity (17), and plays a role in terminating Ca^2+^-induced Ca^2+^ release (18,19), an inherently self-sustaining mechanism which, if unchecked, may deplete intracellular Ca^2+^ stores (20). Sorcin also activates sarco/endoplasmic reticulum Ca^2+^-ATPase (SERCA) pumps and ensures efficient refilling of ER Ca^2+^ stores after contraction (21).

We have recently reported (22) that transgenic mice overexpressing sorcin in the pancreatic beta cell under the rat insulin promoter (RIP7*Sri*TetOn) had improved glucose tolerance and beta cell function when fed a high fat diet (HFD), but had no noticeable phenotype on a standard chow diet. Conversely, mice deleted for the sorcin gene (*Sri*^-/-^) were glucose intolerant and had defective *in vivo* glucose-stimulated insulin secretion (GSIS). Beneficial effects of sorcin in the pancreatic beta cell included increased ER Ca^2+^ stores and glucose-induced intracellular Ca^2+^ fluxes, decreased expression of the islet specific isoform of glucose-6-phosphatase (G6PC2), increased nuclear factor of activated T-cells (NFAT) signalling and an apparent reduction in ER stress. Indeed, sorcin overexpression under lipotoxic conditions prevented the induction of the ER stress markers C/EBP homologue protein (CHOP/DDIT3) and glucose-regulated protein 78/Binding immunoglobulin protein (GRP78/BiP). Interestingly, however, we also reported that sorcin seemed to *increase* the activity of activating transcription factor 6 (ATF6), another marker of ER stress (23), but also a key transcription factor now known to mitigate diabetes risk (24).

Here, in order to investigate further the relationship between sorcin and ATF6, we describe changes in endogenous sorcin expression during ER and lipotoxic stress and the effects of sorcin overexpression or inactivation on ATF6 signalling and ER stress. Our data suggest that sorcin enhances the transcriptional activity of ATF6, and that sorcin dysregulation during the progression of lipotoxicity might compromise ATF6 activity during ER stress, potentially leading to defects in insulin secretion, beta cell mass, and diabetes.

## RESULTS

### ER stress induces sorcin expression

Having previously documented (22) that sorcin protects against lipotoxicity-induced ER Ca^2+^ depletion and pancreatic beta cell dysfunction, we next sought to investigate potential mechanisms behind such protection. We first measured sorcin mRNA levels in response to ER stress-inducing agents, namely thapsigargin and tunicamycin in human islets of Langerhans, the murine insulinoma cell line MIN6, and in the non-excitable human cell line HEK293. CHOP mRNA levels were measured in parallel as a marker of ER stress induction (25).

As shown in Figure 1A, exposure of isolated human islets of Langerhans to tunicamycin and thapsigargin for 24h increased SRI and CHOP mRNA levels significantly (Fig. 1A top panel, SRI fold changes 1.32 ± 0.08 (tunic) and 1.41 ± 0.07 (thaps), p<0.01, n=3; Fig. 1A bottom panel, CHOP fold changes 8.12 ± 0.84 (tunic, p<0.01), and 7.23 ± 0.49 (thaps, p<0.001), n=3). Thapsigargin exposure for 24h also increased SRI and CHOP mRNA levels significantly in MIN6 cells (Fig. 1B top panel, SRI fold change 1.50 ± 0.06 to 1.61 ± 0.02, p<0.005, n=3; Fig. 1B bottom panel, CHOP fold change 11.53 ± 2.03 to 12.31 ± 2.26, p<0.05, n=3) and HEK293 cells (Fig. 1C top panel, SRI fold change 1.30 ± 0.07 to 1.43 ± 0.08, p<0.05, n=2-5; Fig. 1C bottom panel, CHOP fold change 8.44 ± 0.51 to 8.84 ± 0.73, p<0.001, n=3-5).

**Figure 1.**
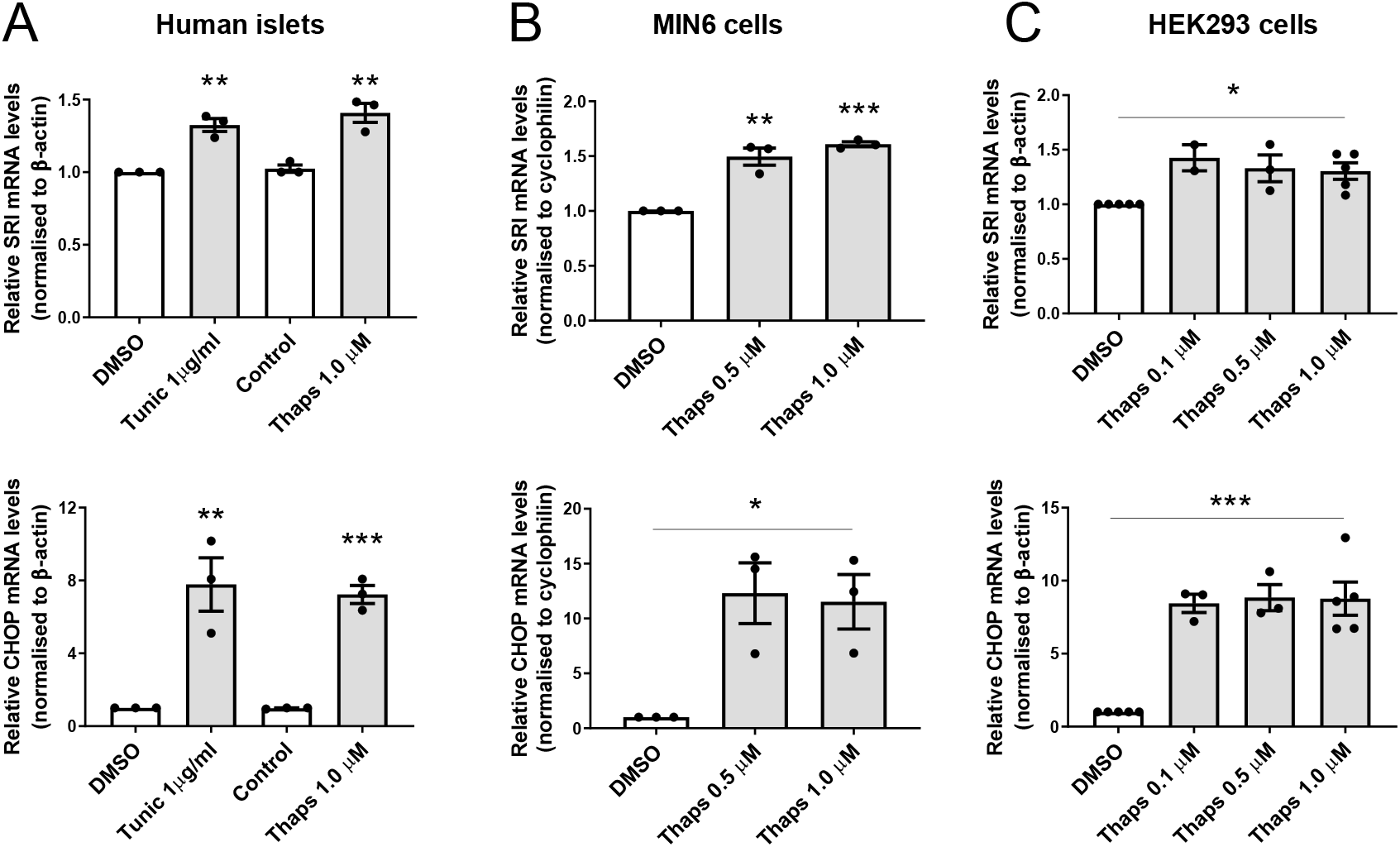
ER stress increases sorcin expression in three different cell types. Human islets (A), MIN6 cells (B) and HEK293 cells (C) were incubated in the presence of DMSO, thapsigargin or tunicamycin as indicated for 24h before qRT-PCR analysis for sorcin (SRI, top panels) and CHOP (lower panels) expression. Results are expressed as means ± S.E.M of ≥ 3 independent experiments; *p<0.05, **p<0.005 and ***p<0.001 by two tailed unpaired Student’s T tests for the effects of ER stress inducers compared to respective controls.

### Palmitate prevents ER stress-induced activation of sorcin expression

In contrast to the above results, induction of ER stress with palmitate in HEK293 cells, failed to increase sorcin mRNA levels despite robust activation of CHOP (Fig. 2A, palmitate-induced CHOP fold change 6.14 ± 0.31, p<0.001, n=4; palmitate-induced sorcin fold change 1.07 ± 0.04, p=NS, n=5). Moreover, prolonged incubation (72h) with palmitate resulted in a decrease in sorcin mRNA levels (Fig. 2B, mRNA sorcin fold change 0.70 ± 0.06, p<0.005, n=5) and protein levels (Fig. 2C, D).

**Figure 2.**
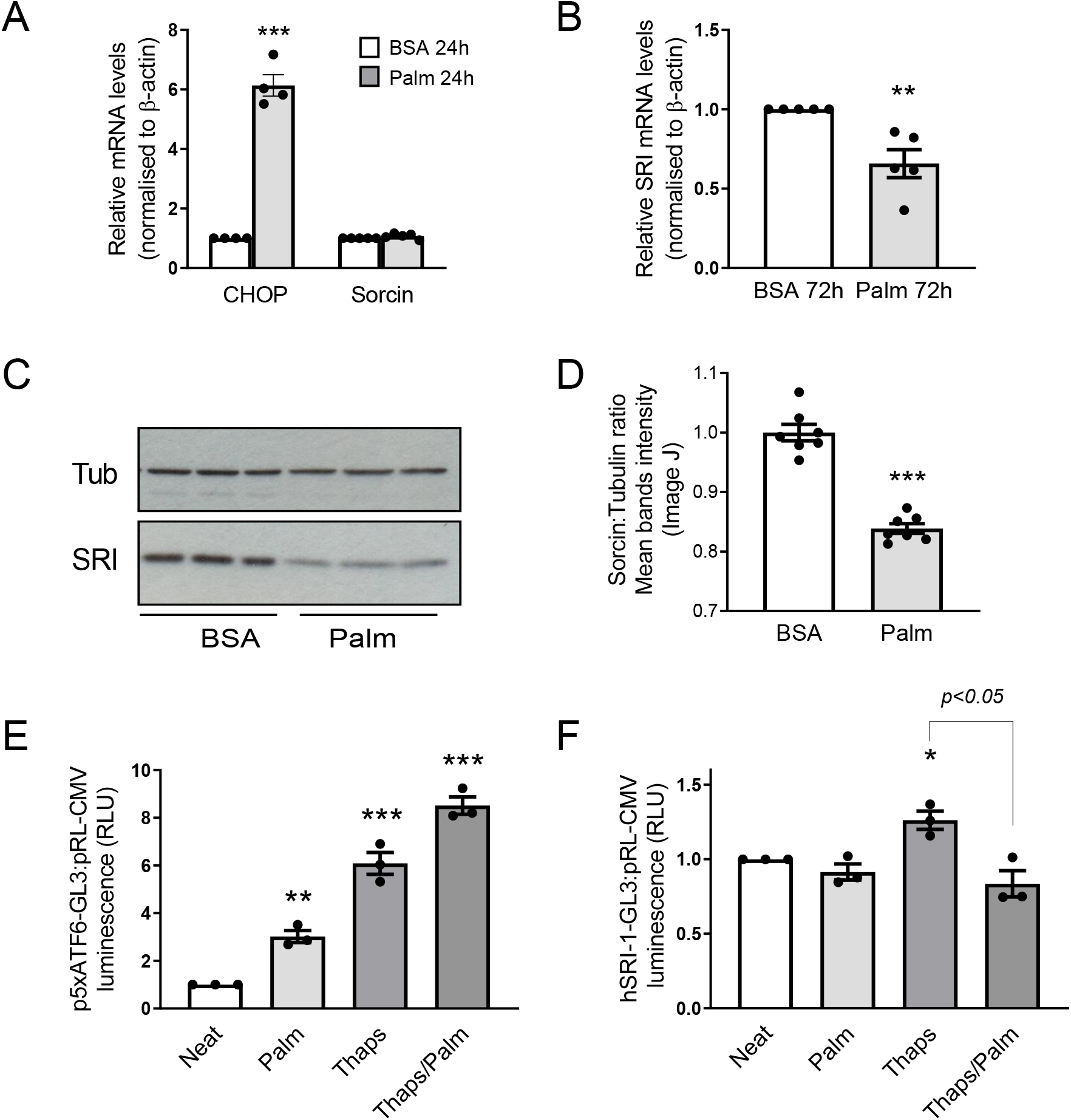
Palmitate prevents ER stress-induced activation of sorcin expression. (A) HEK293 cells were incubated in the presence of 500 μM palmitate for 24h before qRT-PCR analysis for CHOP and sorcin expression. (B) HEK293 cells were incubated in the presence of 500 μM palmitate or BSA vehicle for 72h as indicated before qRT-PCR analysis for sorcin expression. (C, D) Representative Western blot (C) and quantificaton of three independent Western blots (D) of HEK293 cells incubated in the presence of 500 μM palmitate for 72h before total protein extraction and immunoblotting using anti-tubulin and anti-sorcin antibodies as indicated. (E, F) HEK293 cells were transfected with p5xATF6-GL3 (E), or hSRI-1-GL3 (F) and pRL-CMV as transfection standardization control for 24h before medium change containing either palmitate 500 μM, thapsigargin 0.1 μM or a combination of both as indicated for a further 24h before cell lysis and luciferase assay as described in Materials and Methods. Results are expressed as means ± S.E.M of ≥ 3 independent experiments, *p<0.05, **p<0.005 and ***p<0.001 by two tailed unpaired Student’s T tests for the effects of thapsigargin, palmitate or both compared to controls.

To investigate whether the observed changes in sorcin expression were exerted at the transcriptional level, we transfected HEK293 cells with phSRI-1-GL3 or p5xATF6-GL3 for 24h before incubation with thapsigargin, palmitate or a combination of both for a further 24h as indicated in Fig. 2E&F. phSRI-1-GL3 is a *Firefly* luciferase reporter driven by the proximal 1378 bp of the human sorcin promoter whereas p5xATF6-GL3 is a *Firefly* luciferase reporter driven by five tandem repeats of ATF6 binding sites used to monitor the induction of ER stress in parallel.

As expected, p5xATF6-GL3 activity increased proportionally in response to ER stress inducing agents (Fig. 2E): up to 3-fold with palmitate alone (p<0.005, n=3), ~6-fold with thapsigargin alone (p<0.001, n=3) and ~8.5-fold with a palmitate/thapsigargin combination (p<0.001, n=3). However, the human SRI-1 promoter was not activated by palmitate, despite being significantly activated in response to thapsigargin. Remarkably, concomitant incubation with palmitate blocked phSRI-1 promoter activation by thapsigargin (Fig. 2F).

Taken together, these results indicate that lipotoxicity impairs ER stress-induced sorcin expression at the transcriptional level.

### Sorcin enhances ATF6 signalling

ATF6 is a large 90 kDa protein anchored at the ER membrane, which translocates to the Golgi apparatus in response to ER stress. Once in the Golgi, it is cleaved to release a smaller 50 kDa active protein, which enters the nucleus and acts as a transcription factor (23). Therefore, to assay ATF6 signalling, we used two different ATF6 reporter luciferase plasmids, namely p5xATF6-GL3 (26), which contains five tandem repeats of ATF6 binding site and can be activated by endogenous or overexpressed ATF6, and p5xGAL4-Elb-GL3 (23), a GAL4 reporter gene which is exclusively activated by co-transfection with a GAL4-ATF6 fusion protein. The latter was used in order to circumvent any possible non-specific binding on p5xATF6-GL3 reporter by other transcription factors (27), and to specifically measure ATF6 activity in response to sorcin.

As shown in Fig. 3, co-transfection with sorcin increased the activity of p5xATF6-GL3 1.8 fold (p<0.001, n=6, Fig. 3A) and p5xGAL4-Elb-GL3 7.2 fold (p<0.001, n=4-5, Fig. 3B) compared to cotransfection with GFP, used as an inert control protein. Conversely, in *SRI*-null HEK293 cells generated by CRISPR/Cas9-mediated genome editing (see suppl. Fig.1A, B), the activity of both p5xATF6-GL3 and p5xGAL4-Elb-GL3 reporters were reduced by 28% and 26%, respectively, compared to wild type (WT) HEK293 cells (p<0.001, n≥3, Fig. 3C, D). A similar result was obtained in mouse embryonic fibroblast (MEF) cells prepared from sorcin knockout (KO) embryos, where the activity of p5xATF6-GL3 reporter was decreased by 57% (p<0.001, n=5) in basal conditions and by 61% (p<0.05, n=5) in cells stimulated by thapsigargin, compared to WT MEFs (Fig. 3E).

**Figure 3.**
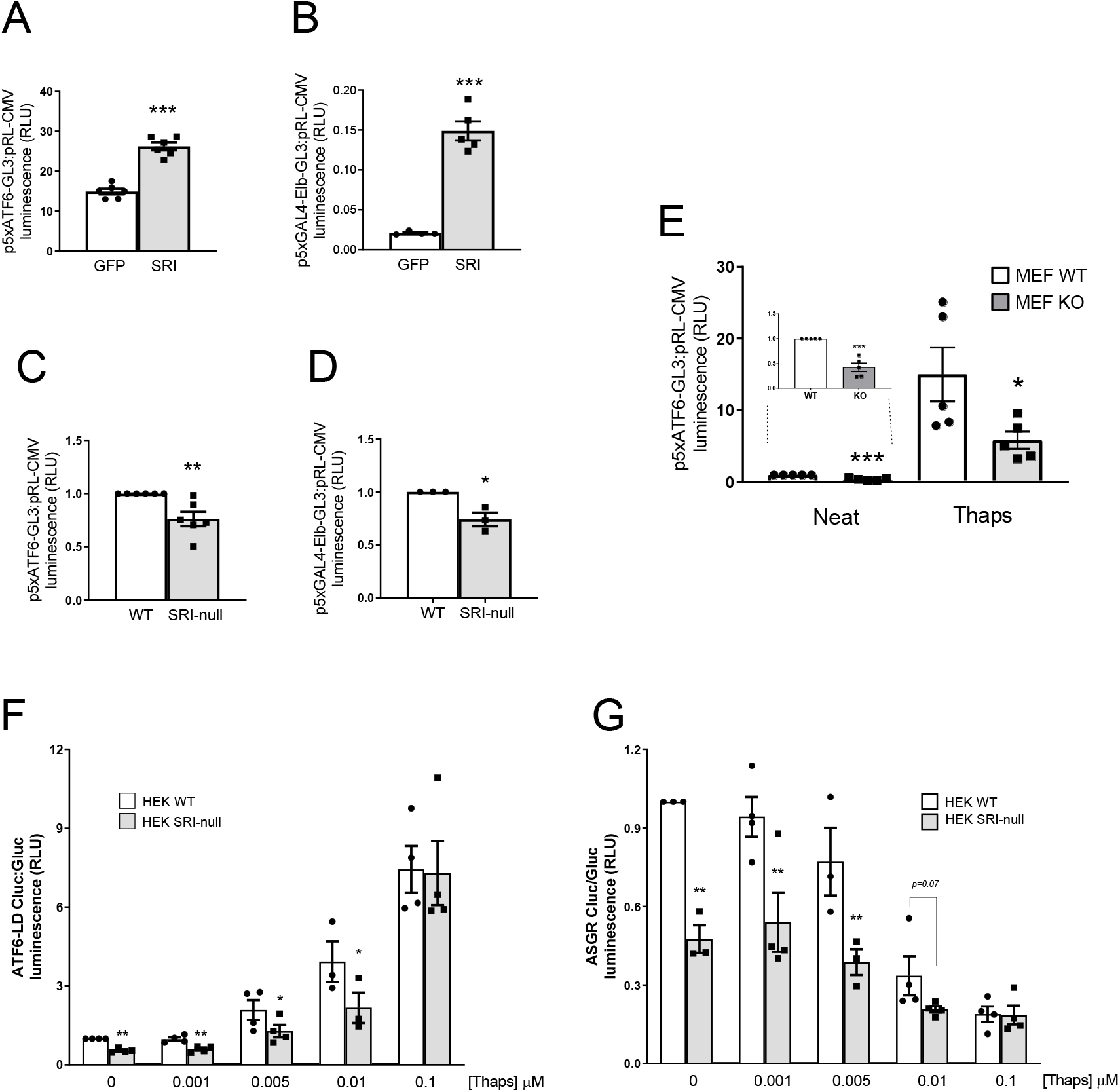
Sorcin stimulates ATF6 activity despite lowering ER stress. (A) HEK293 cells were co-transfected with p5xATF6-GL3 reporter plasmid and either pEGFP-C1 or pSRIm as indicated and (B) HEK293 cells were co-transfected with p5xGAL4-Elb-GL3 and pHA-GAL4-ATF6 and either pEGFP-C1 or pSRIm as indicated for 48h before cell lysis and luciferase assays. (C) WT and SRI-null HEK293 cells were transfected with p5xATF6-GL3 or (D) p5xGAL4-Elb-GL3 and pHA-GAL4-ATF6 for 48h before cell lysis and luciferase assays. (E) SRI WT and KO MEFs were transfected with p5xATF6-GL3 reporter plasmid as in (C) for 24h before medium change containing +/− thapsigargin 0.1 μM as indicated for a further 24h before cell lysis and luciferase assay. (F, G) WT and SRI-null HEK293 cells were transfected with (F) pGluc/ATF6LD-CLuc and (G) pGluc/ASGR-CLuc for 24h before medium change containing increasing doses of thapsigargin as indicated for a further 24h before Gaussia and Cypridina luciferase assays from cell culture supernatants as described in methods. Results are expressed as means ± S.E.M of ≥ 3 independent experiments, *p<0.05, **p<0.005 and ***p<0.001 by two tailed unpaired Student’s T tests.

These data indicate that sorcin enhances ATF6 signalling.

### Sorcin ablation reduces ATF6 activity despite increasing ER stress

We next used a combination of plasmids to measure both specific ATF6 activation and overall cellular ER stress, independently, as described by Fu et al (28). pGLuc/ATF6LD-CLuc encodes *Cypridina* luciferase (CLuc), a secreted luciferase, fused to the C-terminal luminal (intra ER) domain of ATF6, and pGLuc/ASGR-CLuc encodes *Cypridina* luciferase fused to Asialoglycoprotein receptor 1 (ASGR1), a slow maturing protein whose secretion is decreased in ER stress conditions (29). Both plasmids also contain *Gaussia* luciferase (GLuc), another secreted luciferase, under a distinct CMV promoter, to normalise for transfection efficiency. Usually, secretion of both ATF6LD-CLuc and ASGR-CLuc are inversely correlated, as ATF6LD-CLuc secretion increases under conditions of ER stress, whereas ASGR-CLuc secretion decreases (28). We hypothesised that, in the absence of sorcin, both proteins would be less secreted, as this should increase ER stress but also dampen ATF6 activation. Indeed, as shown in Fig. 3F&G, ATF6LD-CLuc and ASGR-CLuc secretion were both reduced by 45% and 52% respectively in *SRI*-null HEK293 cells compared to WT HEK cells in basal conditions. Moreover, this reduction in ATF6 and ASGR secretion in *SRI*-null HEK293 persisted following the addition of low doses of thapsigargin (up to 0.01 μM) but was lost at higher dose of thapsigargin (0.1μM).

Having shown that sorcin was necessary for full ATF6 activity using a variety of luciferase reporters, we next measured expression of endogenous ATF6 target genes in *SRI*-null HEK293 cells in response to ER stress. As shown in Figure 4, the induction of ATF6 target genes GRP78, HERPUD1, SEL1L and EDEM1 was reduced in *SRI*-null cells in response to thapsigargin. On the contrary, sorcin’s absence did not affect induction of CHOP and ATF6 itself by thapsigargin.

**Figure 4.**
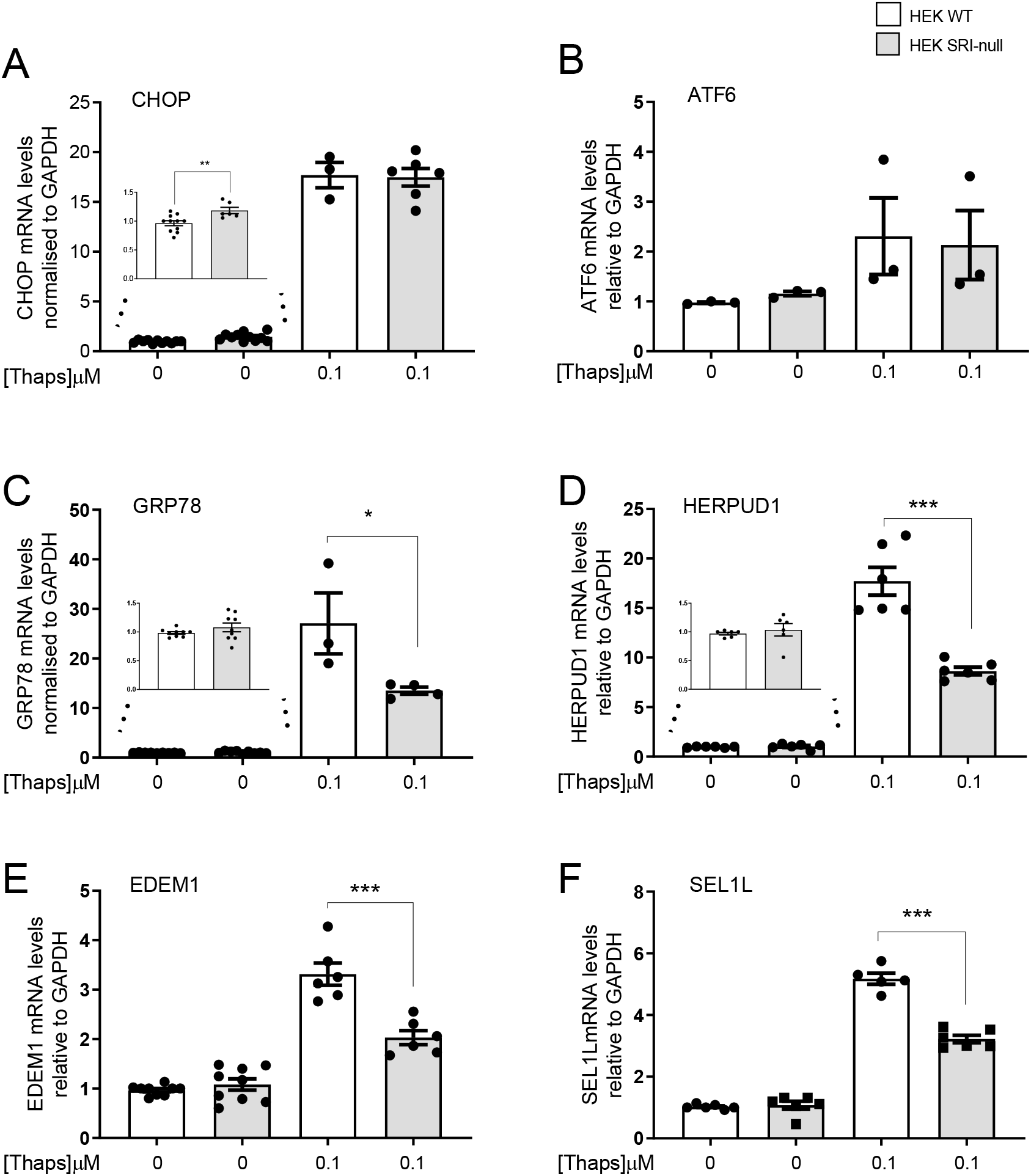
Sorcin inactivation impairs activation of ATF6 target genes during ER stress. (A-F) WT and SRI-null HEK293 cells were incubated with or without 0.1 μM thapsigargin as indicated for 24h before cell lysis and qRT-PCR for CHOP (A), ATF6 (B) and four ATF6 target genes, namely GRP78 (C), HERPUD1 (D), EDEM1 (E), and SEL1L (F) as indicated. Results are expressed as means ± S.E.M of ≥ 3 independent experiments, *p<0.05, **p<0.005 and ***p<0.001 by two tailed unpaired Student’s T tests.

These data confirm that sorcin depletion increases ER stress and that sorcin is necessary for full ATF6 activity.

### Sorcin increases ATF6protein levels

We next proceeded to decipher the mechanism(s) behind sorcin’s regulation of ATF6 activity. Our data above indicated that sorcin influenced both endogenous and exogenous ATF6 protein, since we saw an effect on both p5xATF6-GL3 and p5xGAL4-Elb-GL3 reporters’ activities following manipulation of sorcin protein levels. This suggested a possible effect on ATF6 processing or cleavage (23) rather than ATF6 transcription. Indeed, ATF6 mRNA levels are either slightly elevated or unchanged in *SRI*-null *vs* WT HEK293, probably reflecting the chronic ER stress these cells are under (Fig. 4B and not shown).

To quantify the relative proportion of ATF6p50 (cleaved) to ATF6p90 (full length), we cotransfected HEK293 cells with 3xFLAG-ATF6 together with either sorcin or GFP as control, and treated the cells with thapsigargin or dithiothreitol (DTT), and assayed by western blotting using an anti-FLAG antibody. As shown in Fig. 5A, small (~14%) but reproducible and significant increases were found in the ratios of ATF6p50 band intensity to ATF6p90 band intensity when ATF6 was co-expressed with sorcin compared to GFP in basal condition, although this became less apparent in the presence of high dose thapsigargin (1 μM) and disappeared with DTT.

**Figure 5.**
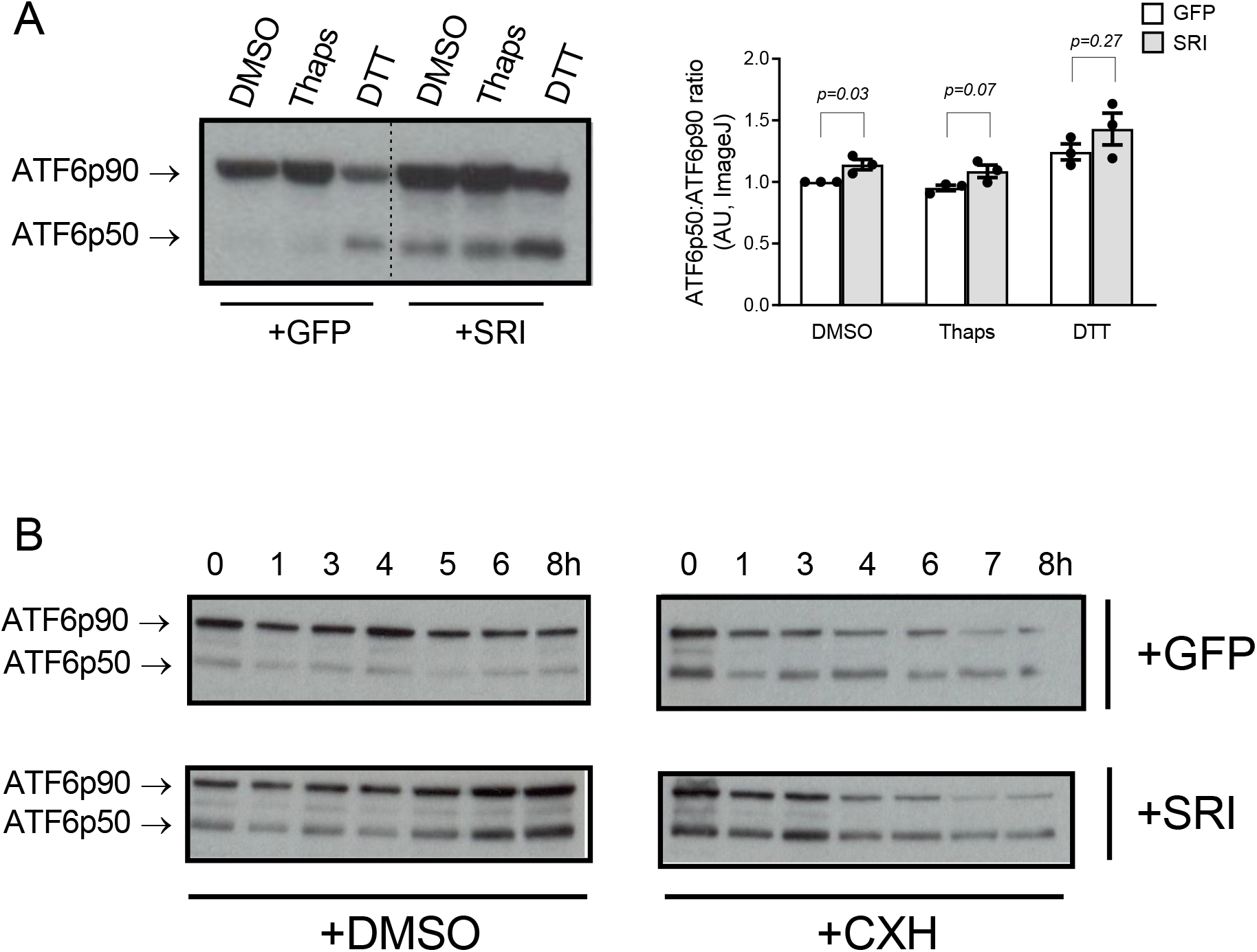
Sorcin increases ATF6 protein levels. (A) HEK cells were transfected with 3xFLAG-ATF6 and either sorcin or GFP as indicated for a total of 48h. Thapsigargin 1 μM and DTT 10 mM were added 16 hours and 1 hour respectively before cell lysis and Western blotting using a monoclonal anti-Flag antibody. Left panel shows a representative Western blot, right panel shows a quantification of 3 independent Western blots by Image J. (B) HEK cells were transfected as above for 24h before the addition of 50 μg/ml cycloheximide (CHX) or DMSO and samples were collected for up to 8h after treatment as indicated. Representative Western blots performed using anti-Flag antibody are shown. Results are expressed as means ± S.E.M of ≥ 3 independent experiments, significance analysis performed by two tailed unpaired Student’s T tests.

However, when sorcin and ATF6 were co-expressed, we invariably observed an increase in total ATF6 immuno-reactivity (not shown). Since it has been reported that ATF6 cleavage is proportional to ATF6 abundance (27,30), we thus sought to investigate whether sorcin might prolong the half-life and/or the stability of ATF6 protein, which in turn would promote its processing and activation. ATF6 protein halflife is short, (2-3 h) (30,31) and ER stress increases ATF6 protein turnover by promoting its degradation through the proteasome (32–34). It was therefore conceivable that, by reducing ER stress (22,35), sorcin might prolong the half-life and/or the stability of ATF6 protein. To test this hypothesis, HEK293 cells were co-transfected with 3xFLAG-ATF6 and sorcin or GFP for 24h and incubated in the presence of cycloheximide (CXH) to stop de novo protein synthesis or DMSO (vehicle) for up to 8 hours before immunoblotting. In control GFP/DMSO conditions (Fig. 5B, top left panel), total ATF6 protein abundance remained fairly constant over the 8 h period, whereas in the presence of overexpressed sorcin (Fig. 5B, bottom left panel), there was a steady increase in the amount of both ATF6p90 and ATF6p50 bands intensity, confirming the positive effect of sorcin on ATF6 abundance and processing. However, sorcin overexpression failed to alter ATF6 immuno-reactivity significantly compared to GFP in the presence of CXH (Fig. 5B, right panels) after 3 hours. This was indicative of increased synthesis rather than decreased degradation.

### Sorcin and ATF6 interact directly during ER stress

We then sought to explore if there could be a direct interaction between sorcin and ATF6 *in cellulo* since Lalioti et al. reported an interaction *in vitro* between the two proteins using a protein array (36). Our repeated attempts to co-immunoprecipitate sorcin (endogenous or overexpressed) and ATF6 using both a Flag-tagged or GFP-tagged ATF6 protein were unsuccessful, either in the presence of 1mM Ca^++^ or 1mM EDTA (not shown). These epitope tags were fused to the cytosolic N-terminal end of ATF6. We next performed Förster/fluorescence resonance energy transfer (FRET) experiments using the acceptor-photobleaching FRET (apFRET) technique (37). In this method, the presence of FRET between a fluorescent protein acting as donor (mCerulean) and a fluorescent protein acting as acceptor (mVenus) is revealed upon inactivation (by photobleaching) of the acceptor. Photobleaching of the fluorescence acceptor attenuates energy transfer between the fluorophores resulting in an increase in donor fluorescence intensity. FRET can only occur over distances between ~ 1-10 nm; thus, an increase in donor fluorescence intensity after photobleaching implies the two fluorophores interact over this distance, suggesting a protein-protein interaction (37).

HEK293 cells were co-transfected with a combination of FRET vectors as illustrated in Fig. 6, including the FRET reference standard C5V (35). mCerulean-tagged ChREBP (Carbohydrate Responsive Element Binding Protein, CFP-ChREBP) was included as a presumed positive control since we previously showed an interaction between ChREBP and sorcin by yeast two-hybrid and co-immunoprecipitation (38). FRET efficiencies between mVenus-tagged sorcin (YFP-SRI) and CFP-ChREBP, CFP-ATF6 (N-terminal mCerulean tag) or ATF6-CFP (C-terminal mCerulean tag) were compared to reference standard vector C5V in basal conditions. FRET efficiencies were also compared in the presence of ionomycin to increase intracellular [Ca^++^] since sorcin conformation changes with intracellular Ca^2+^ concentration and this in turn modifies its interaction with molecular partners (39) and thapsigargin to stimulate ATF6 activity. Negative controls included the pairs CFP-ATF6/YFP alone, CFP alone/YFP-SRI and CFP/YFP.

**Figure 6.**
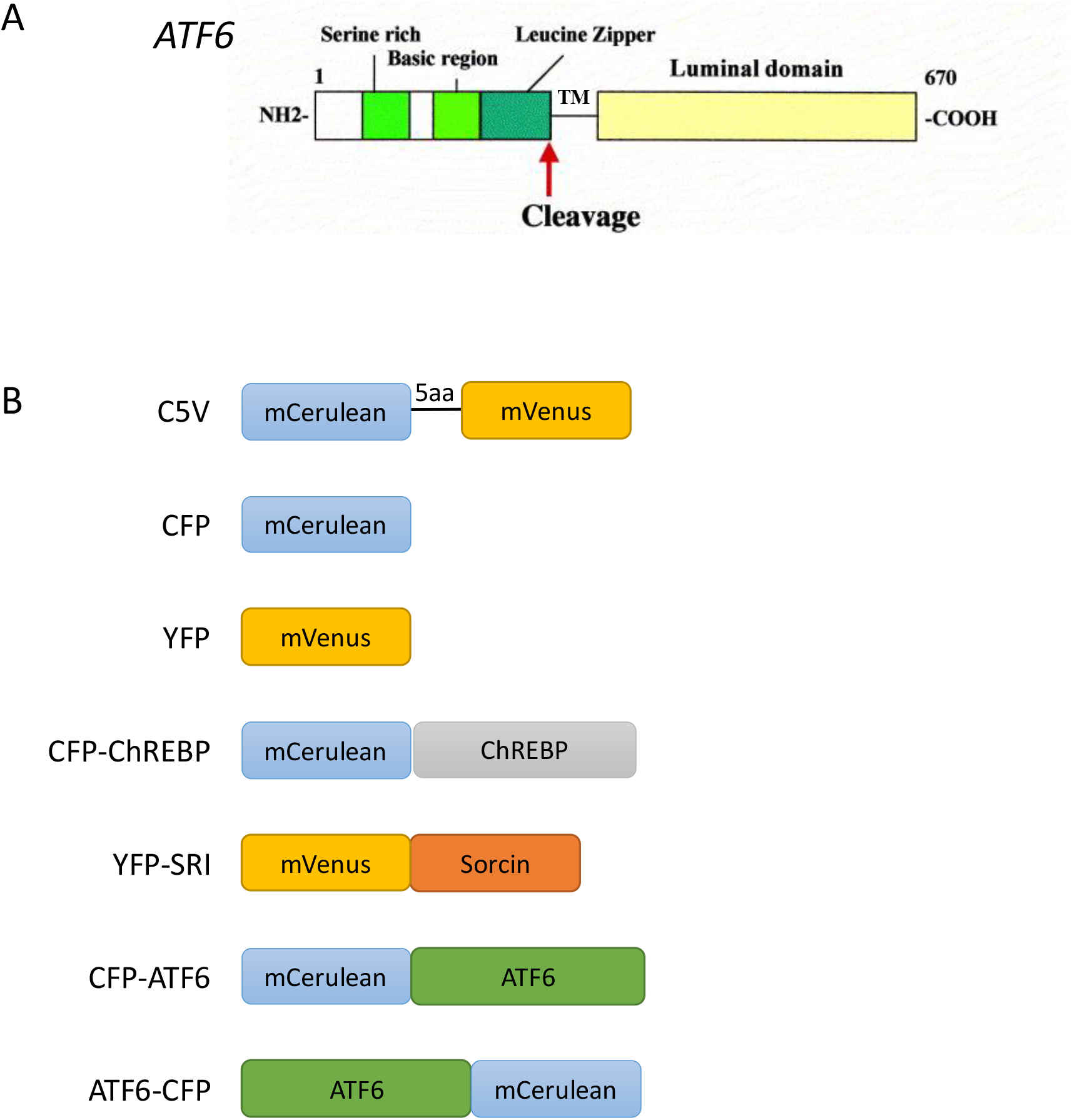
Schematic structures of ATF6 and constructs used in FRET experiments. (A) ATF6 is an ER membrane spanning protein consisting of a C-terminal intra ER domain or luminal domain and a N-terminal cytosolic domain. After translocation to the Golgi following dissociation of the C-terminal domain from BiP/GRP78 in the ER, ATF6 is cleaved by proteases S1P and S2P and the N-terminal domain translocates in the nucleus to become a transcription factor binding ER stress response elements (ERSE) on the promoter of its target genes. Adapted from Imaizumi et al, Biochim Biophys Acta. 2001 May 31;1536(2–3):85-96. (B) Schematic representation of the constructs used in FRET experiments and their corresponding abbreviated names.

As shown in Fig. 7A, in non-treated cells, the FRET efficiency of the reference standard C5V was 24.90 ± 0.95 %, whereas the FRET efficiency between CFP-ChREBP and YFP-SRI was 12.88 ± 1.14 %, significantly lower than C5V but significantly higher than the three negative controls (see below). In comparison, the FRET efficiency of CFP-ATF6/YFP-SRI was lower at 9.03 ± 1.20%, and not significantly higher than the FRET efficiency of the negative controls CFP-ATF6/YFP, CFP/YFP-SRI and CFP/YFP, which were 7.75 ± 1.09 %, 5.30 ± 0.35% and 5.63 ± 0.53 % respectively. Interestingly, the pair ATF6-CFP and YFP-SRI gave a FRET efficiency (11.43 ± 1.30 %) similar to that of CFP-ChREBP and YFP-SRI and was significantly higher than the FRET efficiency of the negative controls.

**Figure 7.**
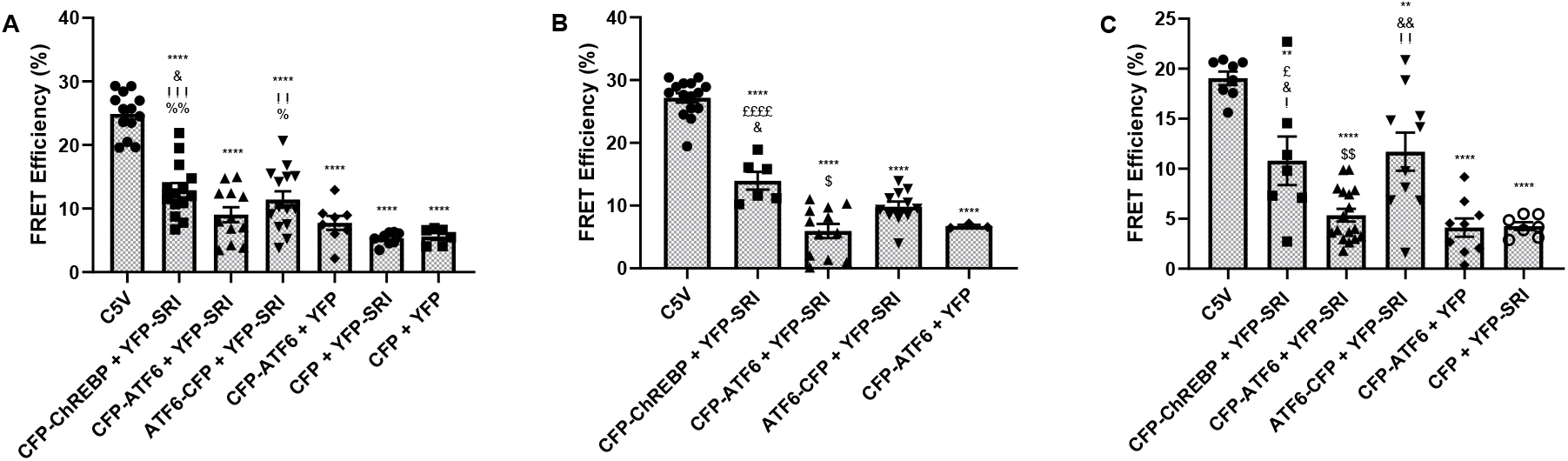
Sorcin and ATF6 Interact at the Protein Level as Measured by Acceptor Photobleaching Förster Resonance Energy Transfer (apFRET). (A-C) Quantification of FRET Efficiency of positive control vector (C5V) and co-transfected Sorcin-mVenus and ATF6-mCerulean plasmids along with appropriate positive and negative controls in (A) non-treated cells, (B) cells treated with ionomycin (1 μM for 1 hour) and (C) cells treated with thapsigargin (0.03μM for 24h). Images were acquired using an LSM 780, AxioObserver confocal microscope and analysed using ImageJ. Mean fluorescence intensity with excitation at 405nm was calculated before and after photobleaching of the chosen regions of interest (ROI) and FRET efficiency calculated as detailed in materials and methods. Results are expressed as mean ± SEM from 3 independent experiments. Significance was calculated using two-tailed Student t test for unpaired data, with Tukey’s multiple comparison test and ANOVA. (*p< 0.05, **p< 0.01, ***p< 0.001, ****p< 0.0001, symbols represent the p-values for the following t-tests: * p-value vs C5V, # vs CFP-ChREBP + YFP-SRI, £ vs CFP-ATF6 + YFP-SRI, § vs ATF6-CFP + YFP-SRI, & vs CFP-ATF6 + YFP, ! vs CFP + YFP-SRI, % vs CFP + YFP)

**Figure 8.**
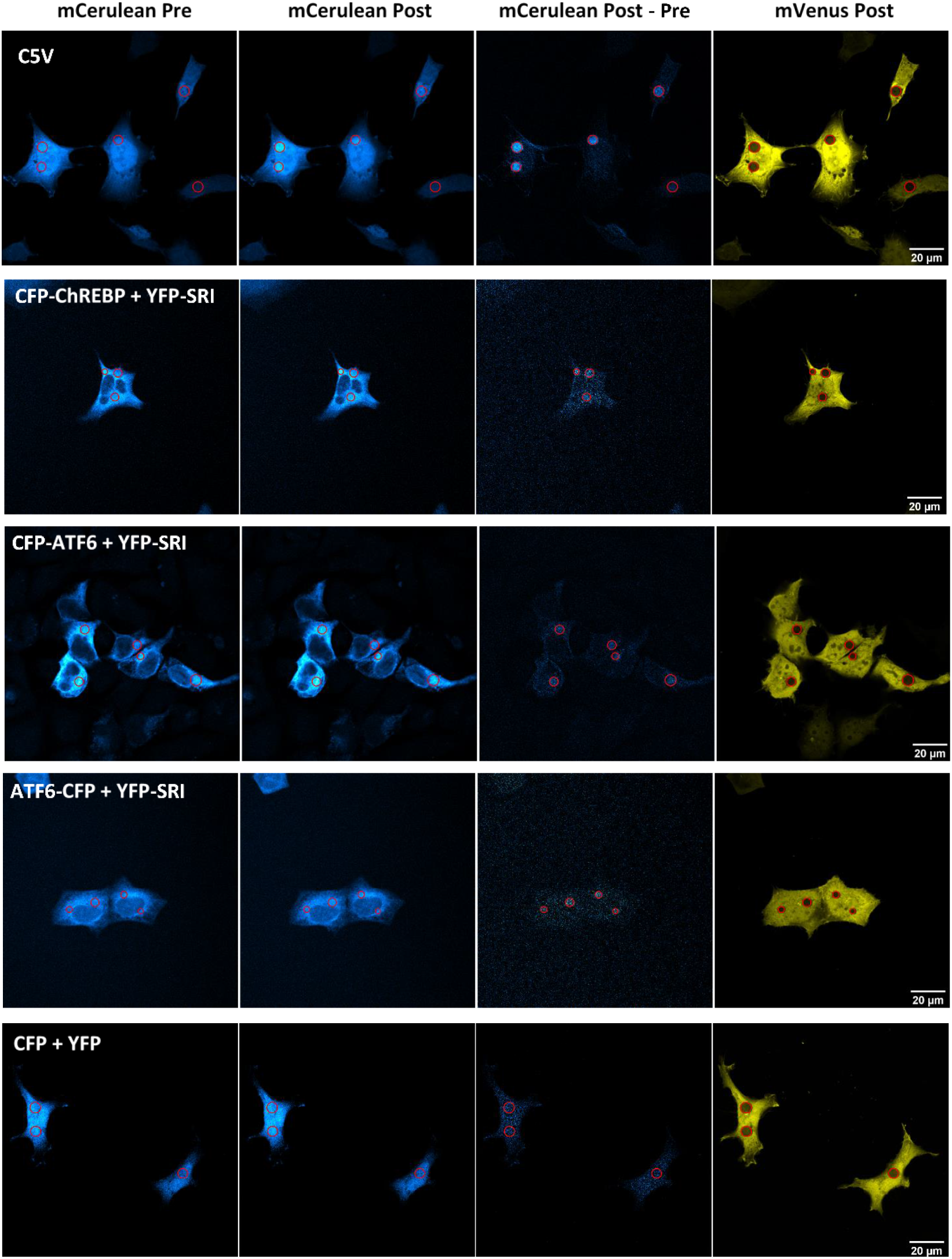
Representative images showing mCerulean fluorescence emission of HEK293 cells transfected with the indicated constructs. Left-Right: mCerulean Pre = mCerulean emission before acceptor photobleaching at 514nm, mCerulean Post = mCerulean emission after acceptor photobleaching at 514nm and mCerulean Post – Pre = net increase in mCerulean emission after photobleaching of mVenus acceptor, calculated using ImageJ software.

In ionomycin-treated cells (Fig 7B; increases in corresponding intracellular [Ca^++^] are shown in suppl. figure 2), the FRET efficiency of C5V was again markedly elevated (27.17 ± 0.76 %) compared to the other combinations of plasmids. FRET efficiency between CFP-ChREBP and YFP-SRI (13.97 ± 1.41 %) was still significantly increased compared to negative control CFP-ATF6 + YFP (6.73 ± 0.21 %). However, FRET efficiency between ATF6-CFP and YFP-SRI (9.83 ± 0.81 %) was no longer significantly different from the negative control CFP-ATF6 + YFP despite being significantly higher than CFP-ATF6 + YFP-SRI combination (5.95 ± 1.13 %).

Finally, in thapsigargin-treated cells (Fig. 7C), the FRET efficiency between the C-terminally tagged ATF6 protein ATF6-CFP and YFP-SRI (11.71 ± 1.91 %) was similar to the FRET efficiency between CFP-ChREBP and sorcin (10.79 ± 2.43 %), but still much lower than the FRET efficiency of C5V (19.04 ± 0.67 %). Again, the FRET efficiency of CFP-ATF6/YFP-SRI was lower at 5.35 ± 0.64 % and not significantly higher than the FRET efficiency of the negative controls CFP-ATF6/YFP and CFP/YFP-SRI at 4.11 ± 0.91 % and 4.26 ± 0.40 % respectively. Representative images for mCerulean emission in the conditions outlined are shown in Fig. 7D, showing the increase in mCerulean emission after photobleaching of the mVenus acceptor in the regions highlighted.

## DISCUSSION

We have presented here a novel molecular explanation for the phenomena known as lipotoxicity (40). The high insulin translation rate makes beta cells extremely sensitive to ER stress (41), which is particularly exacerbated during the progression of obesity (42,43). Sorcin activation during ER stress makes biological sense as an adaptive response, as sorcin has been shown to increase ER Ca^2+^ stores and to decrease apoptosis in response to cytotoxic insults (22,35). However, the saturated fatty acid palmitate, which is the most abundant of the circulating saturated fatty acids (NEFAs) in both obese children (44,45) and adults (46), and associated with beta cell dysfunction (47), decreases sorcin expression both in human islets (22) and in HEK cells (Fig. 2).

Indeed, our data confirm that ER stress induced by thapsigargin and tunicamycin increases sorcin expression (35), but we showed that concomitant exposure to palmitate prevents this beneficial ER stress-induced sorcin transcription. Moreover, longer palmitate exposure reduces sorcin expression (Fig. 2). Thapsigargin blocks the SERCA pump, leading to depletion of ER Ca^2+^ stores and ER stress while tunicamycin inhibits N-linked glycosylation (48), leading to protein misfolding and ER stress. Palmitate also induces ER stress, at least in part by decreasing ER Ca^2+^ stores (3,49). The precise signalling pathway and transcription factor(s) responsible for this palmitate effect on sorcin’s promoter are still unknown but are potentially numerous, including palmitoylation, TNFα, NFκB and Toll-like receptor 4 signalling (50).

Here, we propose another beneficial mechanism of action of sorcin, namely enhancing ATF6 signalling and transcriptional activity.

ATF6 is one of the three ER-resident transmembrane protein sensors that are activated during the UPR, with PKR-like ER kinase (PERK) and Inositol-requiring protein-1 (IRE1). Perturbations in ER homeostasis (accumulation of unfolded proteins, decreased ER Ca^2+^ content, etc.) activates the three branches in parallel to restore ER function. If the UPR is unsuccessful, the cell will undergo apoptosis (51). ATF6 is a type II transmembrane glycoprotein that resides in the ER. Accumulation of misfolded proteins in the ER results in the dissociation of BiP/GRP78 from the C-terminus of ATF6, unmasking a Golgi localisation domain. In the Golgi, ATF6 is processed to its active form through sequential cleavage by site-1 and site-2 proteases (S1P and S2P) (52). The N-terminal cytosolic portion of ATF6 containing a basic leucine zipper motif then translocates to the nucleus and acts as a transcription factor to stimulate the transcription of ER chaperones in order to restore ER homeostasis (30).

ATF6 activation is considered to be the pro-survival branch of the UPR, improving stroke outcomes (53) and myocardial ischemia/reperfusion injury (54,55). In the pancreatic beta cell, ATF6 is needed for beta cell proliferation and compensation during insulin resistance (24) and ATF6-null mice do not display any defects in β-cell function on a normal chow diet but experience severe ER stress with reduced insulin content during HFD (56). These results indicate that ATF6 has an important role in mitigating ER stress and promoting β-cell survival under lipotoxic conditions. The fact that sorcin activates ATF6 signalling is an exciting addition to the known cellular roles of sorcin. Indeed, the beneficial effects of sorcin overexpression *in vivo* in the pancreatic beta cells were only apparent during high fat feeding (22).

Firstly, we have shown that sorcin was necessary for optimum ATF6 transcriptional activity. Using two different *firefly* luciferase reporters, namely p5xATF6-GL3 and p5×GAL4-E1b-GL3, we showed that endogenous and exogenous ATF6 trans-activation ability was stimulated or inhibited following sorcin overexpression or deletion, respectively (Fig. 3A-E). The latter combination of p5×GAL4-E1b-GL3 and pHA-GAL4-ATF6 was used to exclude XBP1s binding or possibly any other non-specific binding activity reported to bind the ERSE of p5xATF6-GL3 (27). Moreover, stimulation of ATF6 target genes during ER stress was also reduced in the absence of sorcin (Fig. 4).

The experiments presented here do provide some explanations about the mechanism(s) behind the stimulatory effect of sorcin on ATF6 signalling. This seems to take place inside the ER through direct interaction between sorcin and the luminal domain of ATF6. Sorcin has been shown to be present in the cytoplasm and the ER by cellular fractionation and to relocate from the ER to the cytosol in response to the antineoplastic agent fluorouracil (FU) which incidentally depletes intra ER Ca^2+^ stores (35).

Indeed, we showed that sorcin ablation decreases the secretion of the fusion protein ATF6LD-CLuc composed of *Cypridina* luciferase (CLuc), a secreted luciferase, fused to the truncated C-terminal luminal (intra ER) domain of ATF6 (Fig. 3F). Also, our FRET experiments have revealed a direct interaction between sorcin-mVenus and ATF6-mCerulean only when the mCerulean tag was fused to the intra-ER C-terminal end of ATF6, but not when it was fused to the cytosolic N-terminal end of ATF6 (Fig. 7).

This was an unexpected result. We initially included ATF6-CFP (which has the fluorescent tag on the C-terminal, intra ER, side of ATF6) as we thought it would be a good negative control but the reverse occurred. Interestingly, ionomycin decreased the FRET efficiency between sorcin and ATF6-CFP whereas thapsigargin seemed to increase it (Fig. 7). As sorcin conformation has been shown to change with intracellular Ca^2+^ concentration (57), and the published *in vitro* interaction between recombinant sorcin and full length ATF6 on a protein array was increased with the addition of 1 mM Ca^2+^ (36), we used the calcium ionophore ionomycin to increase intracellular cytosolic [Ca^2+^] in our FRET experiments, but this seems to decrease the interaction between ATF6-CFP and sorcin (Fig. 7). However, ionomycin has been reported to also decrease ER Ca2+ (58), so more work will be needed to fully understand this interaction. It will be also necessary to confirm the interaction between ATF6-CFP and sorcin by coimmunoprecipitation.

In conclusion, our experiments have shown that sorcin is necessary for full ATF6 activation and transcriptional activity during ER stress, and most likely increases ATF6 protein abundance and signalling through increase protein synthesis and direct interaction with the luminal domain of ATF6.

## EXPERIMENTAL PROCEDURES

### Plasmids generated

Plasmid phSRI-1-GL3, containing the proximal promoter of the human *SRI* gene (transcript variant 1, 198 aa) upstream of luciferase reporter gene, was generated by PCR using human genomic DNA and the following primers: Forward_5’-GAG TTC GGC CCT GAC ATC TA and Reverse_5’-GCA GTC GTC TCC AGC TCT TG. The resulting 1378-bp fragment was sub-cloned into pCR2.1 by TA cloning (Invitrogen), digested by *Kpn*I-*Xho*I and sub-cloned into pGL3basic (Promega) at the same restriction sites.

Plasmid ATF6-CFP, containing human ATF6 cDNA fused to mCerulean on the C-terminal side of ATF6 was generated by amplifying ATF6 ORF from pEGFP-ATF6 (Addgene #32955) by PCR using the following primers: Forward_5’-GTA CTC AGA TCT CGA GCT CAA GC and Reverse_5’ TTG <u>CGG GCC C</u>GT TGT AAT GAC TCA GGG ATG GT *(ApaI* restriction site underlined). The resulting 2046-bp fragment was subsequently sub-cloned into mCeruleanN1 (Addgene #27795) using *HindIII* and *Apa*I restriction sites.

Plasmid YFP-SRI was generated by sub-cloning human sorcin cDNA taken from pAd-hSRI (22) into *KpnI-ApaI* sites of mVenusC1 (Addgene #27794).

Plasmid CFP-ChREBP was generated by sequence and ligation independent cloning (SLIC) (59). cMyc-tagged murine ChREBP ORF was amplified from pChREBP (60) by PCR using the following primers: Forward_5’-<u>GGA CTC AGA TCT CGA</u> GAA CAG AAG CTT ATT TCT GAA GAA G and Reverse_5’ <u>GAA GCT TGA GCT CGA</u> TTA TAA TGG TCT CCC CAG GGT GC (sequences homologous to the destination vector mCeruleanC1 underlined). *Apa*I linearized mCeruleanC1 plasmid and ChREBP PCR product were incubated for 3 minutes in the presence of T4 DNA polymerase before transformation into E. coli to allow for homology directed repair, yielding the CFP-ChREBP construct.

All constructs were verified by DNA sequencing. The other plasmids used in this study are listed in table 1.

**Table 1.** List of other plasmids usedCell culture and transfection

| Plasmid name | Description | Source | Reference |
| --- | --- | --- | --- |
| pSRIm | Expression vector containing murine sorcin cDNA under the control of CMV promoter. | Isabelle Leclerc, Imperial College | (38) |
| p5xATF6-GL3 | *Firefly* luciferase reporter driven by five tandem repeats of ATF6 binding sites. | Ron Prywes, Addgene #11976 | (26) |
| p3xFLAG-ATF6 | Expression vector containing the human ATF6 cDNA with an N-terminal 3xFLAG tag. | Ron Prywes, Addgene #11975 | (23) |
| pHA-GAL4-ATF6 | Expression vector containing an N-terminal HA epitope tag, the N-terminal 147 residues of yeast GAL4 fused to human ATF6 cDNA under the control of SV40 promoter. | Ron Prywes, Columbia University | (23) |
| p5xGAL4-E1b-GL3 | *Firefly* luciferase reporter driven by five GAL4 DNA binding sites. | Ron Prywes, Columbia University | (23) |
| pGluc/ATF6LD-CLuc | Expression vector encoding *Cypridina* luciferase (CLuc), a secreted luciferase, fused to the C-terminal luminal (intra ER) domain of human ATF6 (amino acids 400 to 670). Also contains *Gaussia* luciferase (GLuc), another secreted luciferase, under a distinct CMV promoter, to normalise for transfection efficiency. | Gökhan Hotamisligil, Harvard University | (28) |
| pGluc/ASGR-CLuc | Expression vector encoding *Cypridina* luciferase fused to Asialoglycoprotein receptor 1 (ASGR1), a slow maturing protein whose secretion is decreased in ER stress conditions. Also contains *Gaussia* luciferase (GLuc), another secreted luciferase, under a distinct CMV promoter, to normalise for transfection efficiency. | Gökhan Hotamisligil, Harvard University | (28) |
| CFP-ATF6 | Expression vector containing human ATF6 cDNA with an N-terminal mCerulean tag. | Dhananjaya Kalvakolanu, University of Maryland | (61) |
| C5V | FRET reference standard containing a fusion protein between mCerulean and mVenus separated by 5 amino acids. | Steven Vogel, Addgene #26394 | (62) |
| pLV-CMV-XBP-Gluc | Expression vector containing the first 558 nucleotides of human XBP1 coding sequence (containing the nonconventional XBP1 RNA splicing site by IRE1α) fused to the last 504 nucleotides of GLucM43I, a detergent-resistant variant of the secreted *Gaussia* luciferase. | Arnaud Zaldumbide, Leiden University | (63) |
| pEGFP-C1 | Expression vector containing enhanced Green Fluorescent Protein under the control of CMV promoter. | Clontech® |  |
| pGL3-promoter | Transfection normalisation standard containing *Firefly* luciferase under the control of SV40 promoter. | Promega® |  |
| pRL-CMV | Transfection normalisation standard containing Renilla luciferase under the control of CMV promoter. | Promega® |  |

HEK293 and MIN6 cells were maintained in Dulbecco’s Modified Eagle’s Medium (DMEM) medium as described (22). EndoC-ßH1 cells were maintained on matrigel-fibronectin–coated (100 μg/ml and 2 μg/ml, respectively) culture wells in DMEM that contained 5.6 mM glucose, 2% BSA fraction V (Roche Diagnostics), 50 μM 2-mercaptoethanol, 10 mM nicotinamide (Calbiochem), 5.5 μg/ml transferrin (Sigma-Aldrich), 6.7 ng/ml selenite (Sigma-Aldrich), 100 U/ml penicillin, and 100 μg/ml streptomycin as described in (64). Mouse Embryonic Fibroblasts (MEFs) were prepared from sorcin knockout embryos (*Sri*^-/-^) (22,65) and wild type littermate embryos at E13.5-E15.5 day post-coitum essentially as described in (66) with the omission of DNase I in the digestion step and gelatine in the plating step. Animals were kept at the Imperial College Central Biomedical Service and procedures approved by the UK Home Office Animals Scientific Procedures Act, 1986 (HO License PPL PA03F7F07 to I. L.). MEFs were cultured in DMEM containing 25 mM glucose, 4mmol/l L-glutamine, 100IU/ml penicillin, 100μg/ml streptomycin and 10% (vol/vol) FCS in a humidified atmosphere at 37C with 5% CO_2_. Experiments were performed on MEFs between passages 4 and 20. HEK293 cells were transfected using the calcium phosphate co-precipitation method, pH 7.12, whereas MIN6 cells and MEFs were transfected with either Lipofectamine 2000 or 3000 (Invitrogen) or Xfect (Clontech) as per manufacturer’s instructions. Human islets of Langerhans from normoglycaemic donors were obtained from facilities in Pisa, Oxford and Geneva and cultured as previously described (22).

### CRISPR/Cas9 gene editing

For generating *SRI*-null HEK293 cells, oligonucleotides encompassing guide RNA sequences from the human *SRI* gene located in exon 3 at position 37 (hSRI_Ex3_37Sens: 5’CAC CGC TGA CAC AGT CTG GCA TTG C, hSRI_Ex3_37ASens: 5’AAA CGC AAT GCC AGA CTG TGT CAG) were annealed and phosphorylated before sub-cloning in pX330 at *BsbI* as described in (67). The resulting pX330-hSRI-Ex3-37 plasmid was verified by sequencing before transfection in HEK cells. Twenty-four hours after transfection, the cells were seeded in 96-well plates at a density of ≤1 cell/well for clonal selection. *SRI* gene disruption was confirmed by qRT-PCR and Western blotting on selected clones.

### RNA extraction and qRT-PCR

Total RNA extraction and qRT-PCR were performed as in (68) using the primers listed in Table 2 below.

**Table 2.**
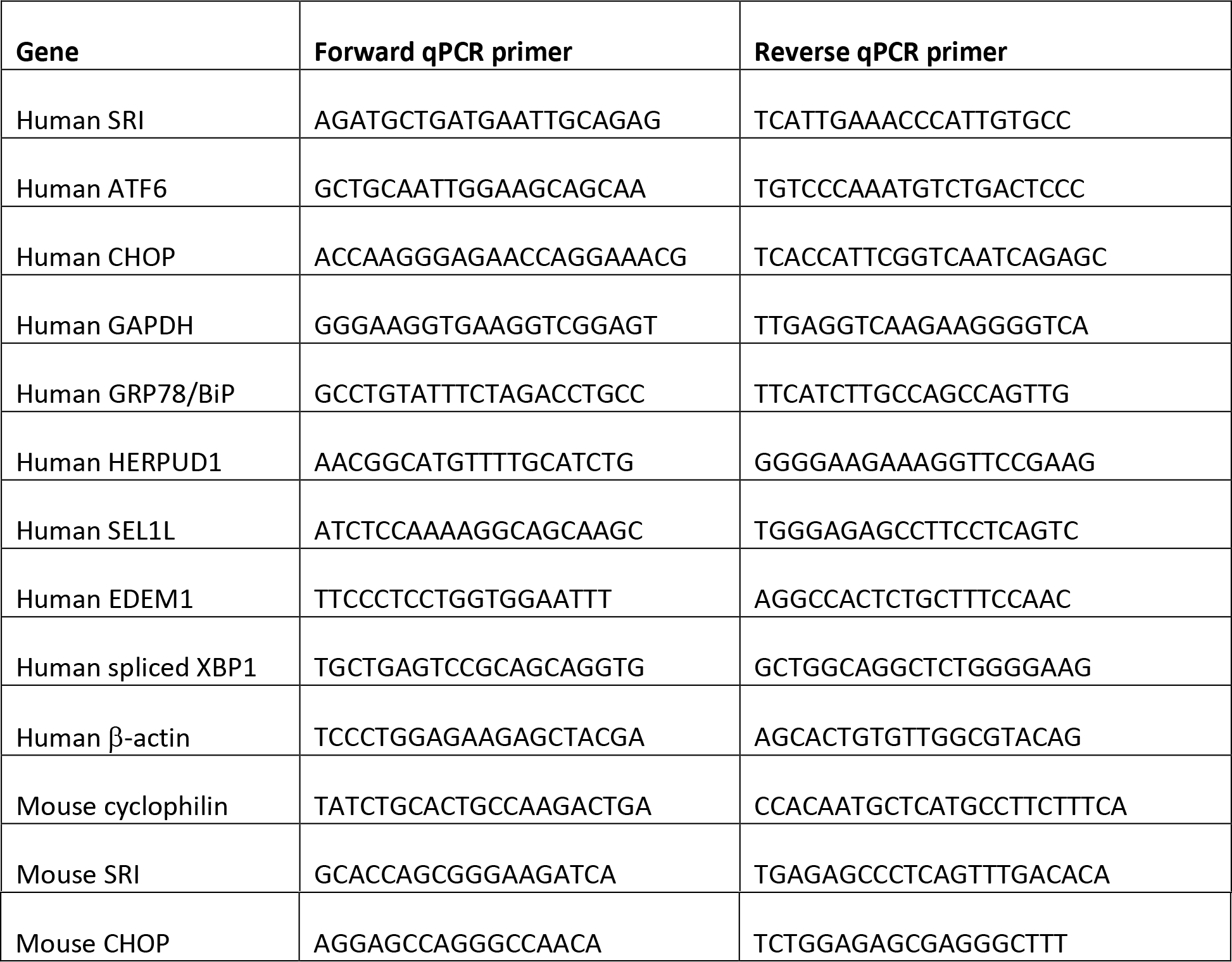
List of qPCR primers

### Western blots

Protein extraction was performed in ice cold RIPA buffer containing 50mM Tris-HCl pH 7.4, 150 mM NaCl, 1% NP40, 0.1% SDS, 0.5% Na Deoxycholate and Roche Complete Proteases Inhibitors. Cells lysates were sonicated for 10 secs and centrifuged at 16,000 *g* for 20 min at 4C before protein concentration assay using Pierce BCA kit. Ten to 25 μg of protein was loaded on 10% acrylamide mini gels, transferred on PVDF membranes and incubated with specific antibodies. Mouse anti-FLAG (Sigma F1804) and mouse anti-tubulin (Sigma T5168) were used at dilution of 1:1000 and 1:10000, respectively. Rabbit anti-sorcin was described previously (19) and used at dilution 1:15000. Quantification of bands intensity was performed using ImageJ software.

### Reporters’ luciferase assays

*Firefly* and *Renilla* luciferases were assayed from whole cell lysates with the Dual-Luciferase^®^ Reporter Assay System (Promega) as per manufacturer’s instructions. *Cypridina* luciferase was assayed from undiluted cell culture supernatants using Pierce^®^ Cypridina Luciferase Glow Assay Kit (Thermo Scientific) or BioLux^®^ Cypridina Luciferase Assay Kit (New England Biolabs) and *Gaussia* luciferase was assayed from 1:40 diluted cell culture supernatant using either the Stop & Glo reagent from the DualLuciferase^®^ Reporter Assay System (Promega) or BioLux^®^ Gaussia Luciferase Assay Kit (New England Biolabs) in a Lumat LB 9507 luminometer (Berthold Technologies).

### FRET imaging studies

Wild-type HEK293T cells were plated on coverslips in 6-well plates and transfected using 2μg of DNA per well by CaPO4 method with the constructs outlined in figure legends. After ~16h, fresh media or media containing 0.03μM thapsigargin was added for a further 24 hours to allow peak expression of the fluorescent proteins. Some cells were then treated with 1 μM ionomycin for one hour as indicated prior to fixation by incubation with 4% PFA for 20 minutes, washing in PBS and distilled water and mounting on slides using ProLong Diamond Antifade Mountant (Invitrogen). Acceptor photobleaching FRET (AP-FRET) was performed on an LSM 780, AxioObserver confocal microscope and images were acquired using a 63x/1.40 Oil DIC Plan-Apochromat objective in a 134.95μm x 134.95μm resolution format. Highlighted regions of interest (ROIs) of ~7um were drawn over the cells expressing both constructs. Fluorophores were stimulated with a 405nm laser and emission split using a CFP and YFP filter and five images were acquired before and after photobleaching to confirm stable fluorescence intensity of mCerulean. The highlighted ROIs were photobleached using a 514nm laser line at 100% intensity for 20 iterations to destroy mVenus acceptor. Images were processed using ImageJ, showing mCerulean emission before and after photobleaching using the cyan hot LUT and mVenus emission after photobleaching.

### Statistical analysis

All results are presented as means ± S.E.M. of at least 3 independent experiments. Differences between means were analysed by two-tailed unpaired Student’s t test or two-way ANOVA as specified. *, ** and *** indicate p < 0.05, 0.005 and 0.001 respectively.

## List of abbreviations

ATF6: activating transcription factor 6;
BiP: binding immunoglobulin protein;
CHOP: C/EBP homologue protein;
ChREBP: Carbohydrate Responsive Element Binding Protein;
DTT: dithiothreitol;
ER: endoplasmic reticulum;
FFA: free fatty acids;
GRP78: glucose-regulated protein 78;
HFD: high fat diet;
MEF: mouse embryonic fibroblast;
NEFA: nonesterified fatty acid;
NFAT: nuclear factor of activated T-cells;
RIP: rat insulin promoter;
RyR: ryanodine receptor;
SERCA: sarco/endoplasmic reticulum Ca^2+^-ATPase;
UPR: unfolded protein response.

## Acknowledgements and Funding

Funding was provided by Imperial College London and grants to I.L. from Diabetes UK (BDA:12/0004535 and BDA:16/0005485) and The Rosetrees Trust (A1063); to G.A.R from the Wellcome Trust (WT098424AIA and 212625/Z/18/Z), the MRC (MR/R022259/1, MR/J0003042/1, MR/L020149/1 and MR/N00275X/1) and Diabetes UK (BDA/15/0005275 and BDA 16/0005485). The authors thank Prof Ron Prywes (Columbia University) and Prof Gökhan Hotamisligil (Harvard University) for providing plasmids and Stephen Rothery of the FILM facility at Imperial College London (http://www.imperial.ac.uk/medicine/facility-for-imaging-by-light-microscopy/) for help with FRET imaging studies.

## Authors’ contributions

SP, TG, NJA, JA, PLC and IL performed experiments and analysed data. SP, PLC, GAR and IL designed experiments. PM, PJ, DB, DVK, HHV, GAR and IL contributed resources. SP, PLC, GAR and IL wrote the manuscript. IL is the guarantor of this work and takes responsibility for the integrity and accuracy of the data presented.

---

**Supplemental Figure 1.**
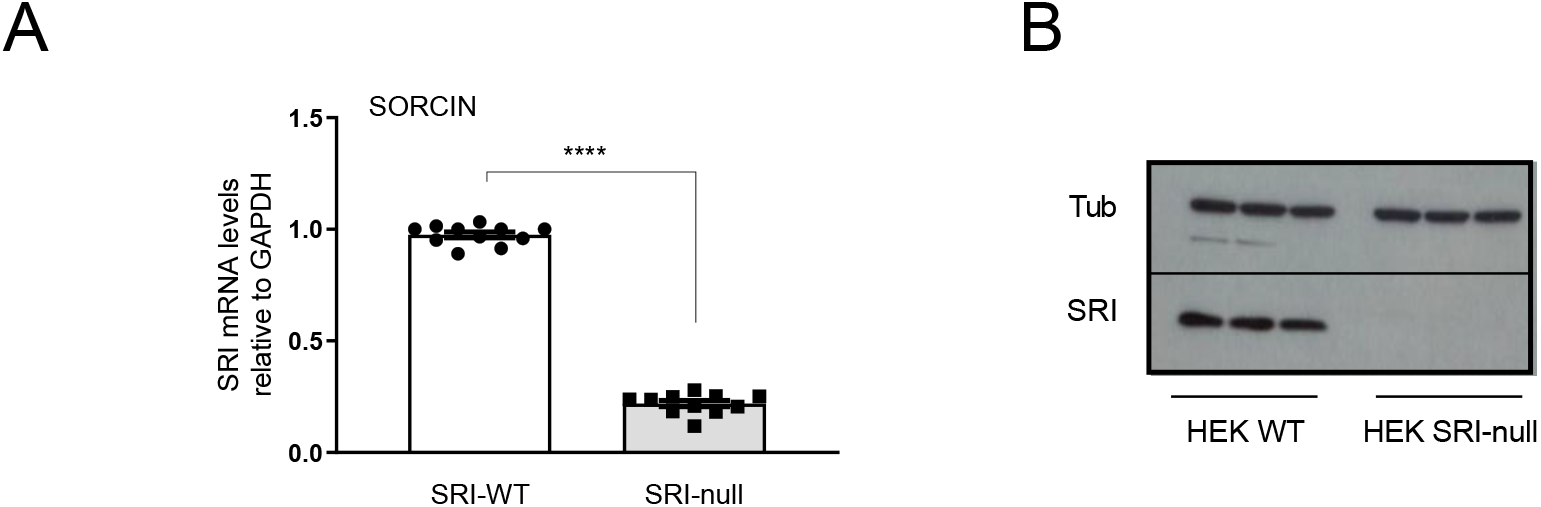
Sorcin mRNA and protein levels in HEK293 WT and HEK293 SRI-null cells. HEK293 SRI-null clonal cells were engeenired by CRISPR/Cas9 gene editing as described in materials and methods before RNA and protein extraction for qRT-PCR (A) and western blot (B) were performed.

**Supplemental Figure 2.**
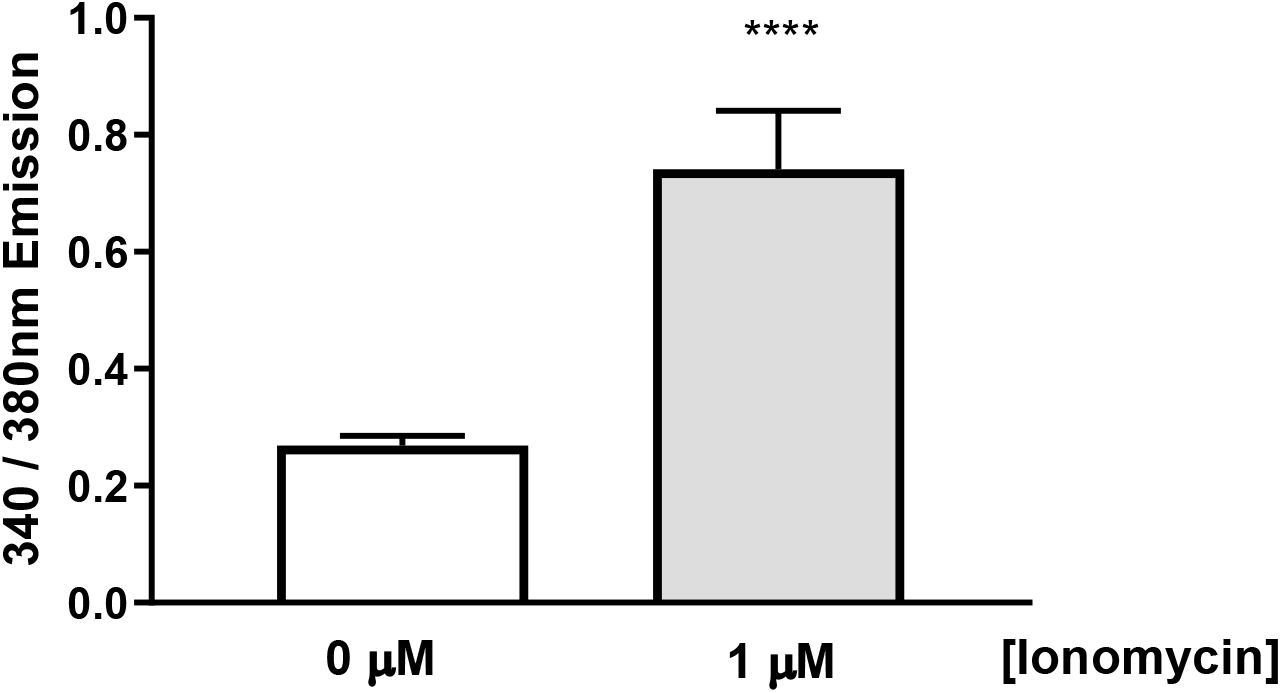
Intracellular Ca^2+^ concentration following inonomycin treatment. HEK293 cells were incubated with Kreb’s buffer containing the trappable intracellular fluorescent Ca^2+^ dye Fura-2-AM. Cells were seeded in 6 well plates at a density of 100,000 cells per well on coverslips and left to recover overnight. Coverslips were then incubated with Kreb’s buffer containing Fura-2-AM (2 μM) with or without ionomycin (1μM) for 1 hour. Following incubation, coverslips were removed from solution and washed before placing in imaging chamber in Kreb’s buffer with or without ionomycin. Emission of individual cells stimulated by 340nm and 380nm laser excitation was imaged and analysis carried out in ImageJ using in-lab Macro. 340nm and 380nm emission was averaged over 180s for each cell to ensure a stable measurement was acquired. Data are means ± SEM. Significance was calcultated using two-tailed Student t test for unpaired data, with Tukey’s multiple comparison test and ANOVA. 0μM = 22 cells, 1μM = 19 cells.

## REFERENCES

1. Prospective Studies, C., Whitlock, G., Lewington, S., Sherliker, P., Clarke, R., Emberson, J., Halsey, J., Qizilbash, N., Collins, R., and Peto, R. (2009) Body-mass index and cause-specific mortality in 900 000 adults: collaborative analyses of 57 prospective studies. Lancet 373, 1083–1096

2. Oyadomari, S., Koizumi, A., Takeda, K., Gotoh, T., Akira, S., Araki, E., and Mori, M. (2002) Targeted disruption of the Chop gene delays endoplasmic reticulum stress-mediated diabetes. J Clin Invest 109, 525–532

3. Cunha, D. A., Hekerman, P., Ladriere, L., Bazarra-Castro, A., Ortis, F., Wakeham, M. C., Moore, F., Rasschaert, J., Cardozo, A. K., Bellomo, E., Overbergh, L., Mathieu, C., Lupi, R., Hai, T., Herchuelz, A., Marchetti, P., Rutter, G. A., Eizirik, D. L., and Cnop, M. (2008) Initiation and execution of lipotoxic ER stress in pancreatic beta-cells. J Cell Sci 121, 2308–2318

4. Ozcan, U., Cao, Q., Yilmaz, E., Lee, A. H., Iwakoshi, N. N., Ozdelen, E., Tuncman, G., Gorgun, C., Glimcher, L. H., and Hotamisligil, G. S. (2004) Endoplasmic reticulum stress links obesity, insulin action, and type 2 diabetes. Science 306, 457–461

5. Boden, G., Duan, X., Homko, C., Molina, E. J., Song, W., Perez, O., Cheung, P., and Merali, S. (2008) Increase in endoplasmic reticulum stress-related proteins and genes in adipose tissue of obese, insulin-resistant individuals. Diabetes 57, 2438–2444

6. Koh, H. J., Toyoda, T., Didesch, M. M., Lee, M. Y., Sleeman, M. W., Kulkarni, R. N., Musi, N., Hirshman, M. F., and Goodyear, L. J. (2013) Tribbles 3 mediates endoplasmic reticulum stress-induced insulin resistance in skeletal muscle. Nature communications 4, 1871

7. Ozcan, L., Ergin, A. S., Lu, A., Chung, J., Sarkar, S., Nie, D., Myers, M. G., Jr., and Ozcan, U. (2009) Endoplasmic reticulum stress plays a central role in development of leptin resistance. Cell Metab 9, 35–51

8. Guerrero-Hernandez, A., Leon-Aparicio, D., Chavez-Reyes, J., Olivares-Reyes, J. A., and DeJesus, S. (2014) Endoplasmic reticulum stress in insulin resistance and diabetes. Cell Calcium 56, 311–322

9. Colotti, G., Poser, E., Fiorillo, A., Genovese, I., Chiarini, V., and Ilari, A. (2014) Sorcin, a calcium binding protein involved in the multidrug resistance mechanisms in cancer cells. Molecules 19, 13976–13989

10. Valdivia, H. H. (1998) Modulation of intracellular Ca2+ levels in the heart by sorcin and FKBP12, two accessory proteins of ryanodine receptors. Trends Pharmacol Sci 19, 479–482

11. Ilari, A., Johnson, K. A., Nastopoulos, V., Verzili, D., Zamparelli, C., Colotti, G., Tsernoglou, D., and Chiancone, E. (2002) The crystal structure of the sorcin calcium binding domain provides a model of Ca2+-dependent processes in the full-length protein. J Mol Biol 317, 447–458

12. Van der Bliek, A. M., Meyers, M. B., Biedler, J. L., Hes, E., and Borst, P. (1986) A 22-kd protein (sorcin/V19) encoded by an amplified gene in multidrug-resistant cells, is homologous to the calcium-binding light chain of calpain. Embo J 5, 3201–3208

13. Maki, M., Kitaura, Y., Satoh, H., Ohkouchi, S., and Shibata, H. (2002) Structures, functions and molecular evolution of the penta-EF-hand Ca2+-binding proteins. Biochim Biophys Acta 1600, 51–60

14. Appelblom, H., Nurmi, J., Soukka, T., Pasternack, M., Penttila, K. E., Lovgren, T., and Niemela, P. (2007) Homogeneous TR-FRET high-throughput screening assay for calcium-dependent multimerization of sorcin. Journal of biomolecular screening 12, 842–848

15. Chen, X., Weber, C., Farrell, E. T., Alvarado, F. J., Zhao, Y. T., Gomez, A. M., and Valdivia, H. H. (2017) Sorcin ablation plus beta-adrenergic stimulation generate an arrhythmogenic substrate in mouse ventricular myocytes. J Mol Cell Cardiol 114, 199–210

16. Meyers, M. B., and Biedler, J. L. (1981) Increased synthesis of a low molecular weight protein in vincristine-resistant cells. Biochem Biophys Res Commun 99, 228–235

17. Lokuta, A. J., Meyers, M. B., Sander, P. R., Fishman, G. I., and Valdivia, H. H. (1997) Modulation of cardiac ryanodine receptors by sorcin. J Biol Chem 272, 25333–25338

18. Fabiato, A. (1983) Calcium-induced release of calcium from the cardiac sarcoplasmic reticulum. The American journal of physiology 245, C1–14

19. Farrell, E. F., Antaramian, A., Rueda, A., Gomez, A. M., and Valdivia, H. H. (2003) Sorcin inhibits calcium release and modulates excitation-contraction coupling in the heart. J Biol Chem 278, 34660–34666

20. Stern, M. D., and Cheng, H. (2004) Putting out the fire: what terminates calcium-induced calcium release in cardiac muscle? Cell Calcium 35, 591–601

21. Matsumoto, T., Hisamatsu, Y., Ohkusa, T., Inoue, N., Sato, T., Suzuki, S., Ikeda, Y., and Matsuzaki, M. (2005) Sorcin interacts with sarcoplasmic reticulum Ca(2+)-ATPase and modulates excitation-contraction coupling in the heart. Basic Res Cardiol 100, 250–262

22. Marmugi, A., Parnis, J., Chen, X., Carmichael, L., Hardy, J., Mannan, N., Marchetti, P., Piemonti, L., Bosco, D., Johnson, P., Shapiro, A. M., Cruciani-Guglielmacci, C., Magnan, C., Ibberson, M., Thorens, B., Valdivia, H. H., Rutter, G. A., and Leclerc, I. (2016) Sorcin links pancreatic beta cell lipotoxicity to ER Ca2+ stores. Diabetes 64, 1009–1021

23. Chen, X., Shen, J., and Prywes, R. (2002) The luminal domain of ATF6 senses endoplasmic reticulum (ER) stress and causes translocation of ATF6 from the ER to the Golgi. J Biol Chem 277, 13045–13052

24. Sharma, R. B., O’Donnell, A. C., Stamateris, R. E., Ha, B., McCloskey, K. M., Reynolds, P. R., Arvan, P., and Alonso, L. C. (2015) Insulin demand regulates beta cell number via the unfolded protein response. J Clin Invest 125, 3831–3846

25. Wang, X. Z., Lawson, B., Brewer, J. W., Zinszner, H., Sanjay, A., Mi, L. J., Boorstein, R., Kreibich, G., Hendershot, L. M., and Ron, D. (1996) Signals from the stressed endoplasmic reticulum induce C/EBP-homologous protein (CHOP/GADD153). Mol Cell Biol 16, 4273–4280

26. Wang, Y., Shen, J., Arenzana, N., Tirasophon, W., Kaufman, R. J., and Prywes, R. (2000) Activation of ATF6 and an ATF6 DNA binding site by the endoplasmic reticulum stress response. J Biol Chem 275, 27013–27020

27. Shen, J., and Prywes, R. (2005) ER stress signaling by regulated proteolysis of ATF6. Methods 35, 382–389

28. Fu, S., Yalcin, A., Lee, G. Y., Li, P., Fan, J., Arruda, A. P., Pers, B. M., Yilmaz, M., Eguchi, K., and Hotamisligil, G. S. (2015) Phenotypic assays identify azoramide as a small-molecule modulator of the unfolded protein response with antidiabetic activity. Science translational medicine 7, 292ra298

29. Fu, S., Yang, L., Li, P., Hofmann, O., Dicker, L., Hide, W., Lin, X., Watkins, S. M., Ivanov, A. R., and Hotamisligil, G. S. (2011) Aberrant lipid metabolism disrupts calcium homeostasis causing liver endoplasmic reticulum stress in obesity. Nature 473, 528–531

30. Haze, K., Yoshida, H., Yanagi, H., Yura, T., and Mori, K. (1999) Mammalian transcription factor ATF6 is synthesized as a transmembrane protein and activated by proteolysis in response to endoplasmic reticulum stress. Mol Biol Cell 10, 3787–3799

31. Teske, B. F., Wek, S. A., Bunpo, P., Cundiff, J. K., McClintick, J. N., Anthony, T. G., and Wek, R. C. (2011) The eIF2 kinase PERK and the integrated stress response facilitate activation of ATF6 during endoplasmic reticulum stress. Mol Biol Cell 22, 4390–4405

32. Hong, M., Li, M., Mao, C., and Lee, A. S. (2004) Endoplasmic reticulum stress triggers an acute proteasome-dependent degradation of ATF6. Journal of cellular biochemistry 92, 723–732

33. Yoshida, H., Uemura, A., and Mori, K. (2009) pXBP1(U), a negative regulator of the unfolded protein response activator pXBP1(S), targets ATF6 but not ATF4 in proteasome-mediated degradation. Cell structure and function 34, 1–10

34. Fonseca, S. G., Ishigaki, S., Oslowski, C. M., Lu, S., Lipson, K. L., Ghosh, R., Hayashi, E., Ishihara, H., Oka, Y., Permutt, M. A., and Urano, F. (2010) Wolfram syndrome 1 gene negatively regulates ER stress signaling in rodent and human cells. J Clin Invest 120, 744–755

35. Maddalena, F., Laudiero, G., Piscazzi, A., Secondo, A., Scorziello, A., Lombardi, V., Matassa, D. S., Fersini, A., Neri, V., Esposito, F., and Landriscina, M. (2011) Sorcin induces a drug-resistant phenotype in human colorectal cancer by modulating Ca(2+) homeostasis. Cancer Res 71, 7659–7669

36. Lalioti, V. S., Ilari, A., O’Connell, D. J., Poser, E., Sandoval, I. V., and Colotti, G. (2014) Sorcin links calcium signaling to vesicle trafficking, regulates Polo-like kinase 1 and is necessary for mitosis. PLoS One 9, e85438

37. Snapp, E. L., and Hegde, R. S. (2006) Rational design and evaluation of FRET experiments to measure protein proximities in cells. Current protocols in cell biology / editorial board, Juan S. Bonifacino … [et al.] Chapter 17, Unit 17 19

38. Noordeen, N. A., Meur, G., Rutter, G. A., and Leclerc, I. (2012) Glucose-induced nuclear shuttling of ChREBP is mediated by sorcin and Ca(2+) ions in pancreatic beta-cells. Diabetes 61, 574–585

39. Meyers, M. B., Pickel, V. M., Sheu, S. S., Sharma, V. K., Scotto, K. W., and Fishman, G. I. (1995) Association of sorcin with the cardiac ryanodine receptor. J Biol Chem 270, 26411–26418

40. Poitout, V., and Robertson, R. P. (2008) Glucolipotoxicity: fuel excess and beta-cell dysfunction. Endocr Rev 29, 351–366

41. Scheuner, D., and Kaufman, R. J. (2008) The unfolded protein response: a pathway that links insulin demand with beta-cell failure and diabetes. Endocr Rev 29, 317–333

42. Biden, T. J., Boslem, E., Chu, K. Y., and Sue, N. (2014) Lipotoxic endoplasmic reticulum stress, beta cell failure, and type 2 diabetes mellitus. Trends Endocrinol Metab 25, 389–398

43. Volchuk, A., and Ron, D. (2010) The endoplasmic reticulum stress response in the pancreatic betacell. Diabetes Obes Metab 12 Suppl 2, 48–57

44. Sabin, M. A., De Hora, M., Holly, J. M., Hunt, L. P., Ford, A. L., Williams, S. R., Baker, J. S., Retallick, C. J., Crowne, E. C., and Shield, J. P. (2007) Fasting nonesterified fatty acid profiles in childhood and their relationship with adiposity, insulin sensitivity, and lipid levels. Pediatrics 120, e1426–1433

45. Staaf, J., Ubhayasekera, S. J., Sargsyan, E., Chowdhury, A., Kristinsson, H., Manell, H., Bergquist, J., Forslund, A., and Bergsten, P. (2016) Initial hyperinsulinemia and subsequent beta-cell dysfunction is associated with elevated palmitate levels. Pediatr Res 80, 267–274

46. Carta, G., Murru, E., Banni, S., and Manca, C. (2017) Palmitic Acid: Physiological Role, Metabolism and Nutritional Implications. Front Physiol 8, 902

47. Maris, M., Robert, S., Waelkens, E., Derua, R., Hernangomez, M. H., D’Hertog, W., Cnop, M., Mathieu, C., and Overbergh, L. (2013) Role of the saturated nonesterified fatty acid palmitate in beta cell dysfunction. J Proteome Res 12, 347–362

48. Duksin, D., and Mahoney, W. C. (1982) Relationship of the structure and biological activity of the natural homologues of tunicamycin. J Biol Chem 257, 3105–3109

49. Xu, S., Nam, S. M., Kim, J. H., Das, R., Choi, S. K., Nguyen, T. T., Quan, X., Choi, S. J., Chung, C. H., Lee, E. Y., Lee, I. K., Wiederkehr, A., Wollheim, C. B., Cha, S. K., and Park, K. S. (2015) Palmitate induces ER calcium depletion and apoptosis in mouse podocytes subsequent to mitochondrial oxidative stress. Cell death & disease 6, e1976

50. Fatima, S., Hu, X., Gong, R. H., Huang, C., Chen, M., Wong, H. L. X., Bian, Z., and Kwan, H. Y. (2019) Palmitic acid is an intracellular signaling molecule involved in disease development. Cellular and molecular life sciences: CMLS 76, 2547–2557

51. Szegezdi, E., Logue, S. E., Gorman, A. M., and Samali, A. (2006) Mediators of endoplasmic reticulum stress-induced apoptosis. EMBO Rep 7, 880–885

52. Ye, J., Rawson, R. B., Komuro, R., Chen, X., Dave, U. P., Prywes, R., Brown, M. S., and Goldstein, J. L. (2000) ER stress induces cleavage of membrane-bound ATF6 by the same proteases that process SREBPs. Mol Cell 6, 1355–1364

53. Yu, Z., Sheng, H., Liu, S., Zhao, S., Glembotski, C. C., Warner, D. S., Paschen, W., and Yang, W. (2017) Activation of the ATF6 branch of the unfolded protein response in neurons improves stroke outcome. Journal of cerebral blood flow and metabolism: official journal of the International Society of Cerebral Blood Flow and Metabolism 37, 1069–1079

54. Jin, J. K., Blackwood, E. A., Azizi, K., Thuerauf, D. J., Fahem, A. G., Hofmann, C., Kaufman, R. J., Doroudgar, S., and Glembotski, C. C. (2017) ATF6 Decreases Myocardial Ischemia/Reperfusion Damage and Links ER Stress and Oxidative Stress Signaling Pathways in the Heart. Circ Res 120, 862–875

55. Blackwood, E. A., Azizi, K., Thuerauf, D. J., Paxman, R. J., Plate, L., Kelly, J. W., Wiseman, R. L., and Glembotski, C. C. (2019) Pharmacologic ATF6 activation confers global protection in widespread disease models by reprograming cellular proteostasis. Nature communications 10, 187

56. Usui, M., Yamaguchi, S., Tanji, Y., Tominaga, R., Ishigaki, Y., Fukumoto, M., Katagiri, H., Mori, K., Oka, Y., and Ishihara, H. (2012) Atf6alpha-null mice are glucose intolerant due to pancreatic betacell failure on a high-fat diet but partially resistant to diet-induced insulin resistance. Metabolism 61, 1118–1128

57. Meyers, M. B., Zamparelli, C., Verzili, D., Dicker, A. P., Blanck, T. J., and Chiancone, E. (1995) Calcium-dependent translocation of sorcin to membranes: functional relevance in contractile tissue. FEBS Lett 357, 230–234

58. Bird, G. S., DeHaven, W. I., Smyth, J. T., and Putney, J. W., Jr. (2008) Methods for studying store-operated calcium entry. Methods 46, 204–212

59. Li, M. Z., and Elledge, S. J. (2007) Harnessing homologous recombination in vitro to generate recombinant DNA via SLIC. Nat Methods 4, 251–256

60. da Silva Xavier, G., Rutter, G. A., Diraison, F., Andreolas, C., and Leclerc, I. (2006) ChREBP binding to fatty acid synthase and L-type pyruvate kinase genes is stimulated by glucose in pancreatic beta-cells. J Lipid Res 47, 2482–2491

61. Gade, P., Ramachandran, G., Maachani, U. B., Rizzo, M. A., Okada, T., Prywes, R., Cross, A. S., Mori, K., and Kalvakolanu, D. V. (2012) An IFN-gamma-stimulated ATF6-C/EBP-beta-signaling pathway critical for the expression of Death Associated Protein Kinase 1 and induction of autophagy. P Natl Acad Sci USA 109, 10316–10321

62. Koushik, S. V., Chen, H., Thaler, C., Puhl, H. L., and Vogel, S. S. (2006) Cerulean, Venus, and Venus(Y67C) FRET reference standards. Biophysical Journal 91, L99–L101

63. Kracht, M. J. L., de Koning, E. J. P., Hoeben, R. C., Roep, B. O., and Zaldumbide, A. (2018) Bioluminescent reporter assay for monitoring ER stress in human beta cells. Scientific reports 8, 17738

64. Ravassard, P., Hazhouz, Y., Pechberty, S., Bricout-Neveu, E., Armanet, M., Czernichow, P., and Scharfmann, R. (2011) A genetically engineered human pancreatic beta cell line exhibiting glucose-inducible insulin secretion. J Clin Invest 121, 3589–3597

65. Chen, X., Weber, C., Farrell, E. T., Alvarado, F. J., Zhao, Y. T., Gomez, A. M., and Valdivia, H. H. (2018) Sorcin ablation plus beta-adrenergic stimulation generate an arrhythmogenic substrate in mouse ventricular myocytes. J Mol Cell Cardiol 114, 199–210

66. Jozefczuk, J., Drews, K., and Adjaye, J. (2012) Preparation of mouse embryonic fibroblast cells suitable for culturing human embryonic and induced pluripotent stem cells. Journal of visualized experiments: JoVE

67. Ran, F. A., Hsu, P. D., Wright, J., Agarwala, V., Scott, D. A., and Zhang, F. (2013) Genome engineering using the CRISPR-Cas9 system. Nat Protoc 8, 2281–2308

68. Sun, G., Tarasov, A. I., McGinty, J. A., French, P. M., McDonald, A., Leclerc, I., and Rutter, G. A. (2010) LKB1 deletion with the RIP2.Cre transgene modifies pancreatic beta-cell morphology and enhances insulin secretion in vivo. Am J Physiol Endocrinol Metab 298, E1261–1273

